# Differential Binding Kinetics for Evaluating Immune Checkpoint Inhibitors in Serum

**DOI:** 10.1101/2021.08.29.458058

**Authors:** Danfeng Yao, Heng Yu, Aaron B. Kantor, Sebastian J. Osterfeld, Toshiro Saito, M. Luis Carbonell, Kalidip Choudhury, Shan X. Wang

## Abstract

The ability to characterize the binding kinetics of drug-target interactions in a biologically relevant matrix, such as serum or plasma, remains a fundamental challenge in drug discovery. We apply a novel label-based giant magnetoresistance (GMR) biosensor platform to measure protein binding kinetics and affinities of drug-target pairs in buffer and different levels of serum. Specifically, we evaluate three well-established immune checkpoint inhibitors, pembrolizumab, nivolumab and atezolizumab and compare the results with label-free kinetic platforms: surface plasmon resonance (SPR) and bio-layer interferometry (BLI). Labeling of analytes does not affect their association and dissociation rates (on and off rates) from GMR biosensors which enables kinetic measurements in biologically relevant matrices. Only the GMR biossensors is consistently suitable for measuring binding kinetics in up to 80% serum. The faster and different off-rates of the three immune checkpoint inhibitors in the presence of serum should be considered when modeling their pharmacological performance.

**Teaser:** We reveal the effects of serum on binding kinetics of antibody drugs, relevant to the pharmacological performance of immunotherapeutic.

## Introduction

A comprehensive understanding of the molecular interactions between drugs and target molecules is essential to the development of effective medicines and optimization of the therapeutic index.^1^ Binding kinetics can help differentiate therapeutic properties for drugs that bind the same target. Examples are given in **Supplementary Table 1**.

Systematic evaluation of the interactions between drugs and their target biomolecules often lacks consideration of the myriad of components in the physiologically relevant matrix. Measuring binding kinetics in a relevant matrix is necessary to fully validate the reliability and predictability of *in vitro* assays. Endogenous molecules in blood can exhibit specific and non-specific interactions with a drug and directly determine the pharmacological response.^2^ Studies have suggested that the interactions with blood proteins modulate the efficacy of protein binding by affecting conformational changes in the target protein.^3^ In some extreme situations, antagonist drugs lose their capacity to bind to their receptors. The strengths with which blood proteins bind to their target molecule alters the drug potency, dosing regimen and the degree of off-target side effects. However, the surface plasmon resonance (SPR)^4^ and bio-layer interferometry (BLI)^5^ platforms typically used to measure binding kinetics are unable to consistently and reliably measure kinetics in physiologically relevant matrices. Therefore, critical matrix studies are often limited and use an equilibrium method such as ultrafiltration or equilibrium dialysis, which do not provide binding kinetics information.

The giant magnetoresistive sensor (GMR) platform has been previously described and shown to be suitable for ligand binding assays in biological matrices.^6–9^ In this method, one molecule (capture) is fixed on the GMR sensor and another molecule (analyte) is labeled with magnetic nanoparticles (MNP). As the MNP-analyte complex binds to the capture surface, the magnetic field of the MNP causes a change in the electrical resistances of the underlying GMR sensor. The change in magnetoresistance ratio (ΔMR) expressed in parts per million (PPM), enables the quantitative real-time monitoring of the binding events. Only MNP-labeled molecules near the binding surface are detected. Direct binding from matrix components that are not labeled is not detected. This is a fundamental distinction of GMR compared to SPR and BLI techniques where specific binding and nonspecific binding of serum proteins cannot be distinguished.

We and our colleagues have previously shown that the GMR platform can be used to evaluate the serum matrix effects on the interactions between targets and drug molecules.^10^ Some pairs, like protein kinase A and its small molecule target quercetin, were greatly affected by the presence of a commercial serum-based product. In contrast, the commercial serum-based product had little effect on two antibody drug-target pairs: infliximab-TNFa and bevacizumab-VEGF. Here we greatly expand our GMR studies to measure and contrast the binding kinetics in serum dilutions of three well-established immune checkpoint inhibitors, pembrolizumab, nivolumab and atezolizumab, which have helped to revolutionize the field of onclology.^11–13^ Pembrolizumab and nivolumab bind to programmed cell death protein-1 (PD-1) which is present on the surface of T and B cells. It has two ligands: programmed death-ligand 1 and ligand 2 (PD-L1, PD-L2). Atezolizumab binds to PD-L1 which is commonly expressed on the surface of antigen presenting cells and tumor cells.

We provide robust platform validation and demonstrate clear analytical advantages by comparing the GMR results with two label-free kinetic platforms, SPR and BLI and to equilibrium-based measurements on the GMR platform.^14^ It is noteworthy that the three pairs display different degrees of increased off-rates in the same amount of serum. The serum-buffer difference in k_off_, was the greatest for the atezolizumab/PDL-1 pair, while the difference for the pembrolizumab/PD-1 pair was appreciable greater than for the nivolumab/PD-1 pair. The differential binding kinetics exhibited in serum should be accounted for when the pharmacological kinetics of the drugs are considered.

## Results

### GMR biosensor assay conditions can readily be established

The GMR kinetic assay can be implemented in two orientations, with either the target or the drug as the capture molecule. Here we mostly present results using different levels of target molecules as capture on the GMR chip and directly labeled MNP-conjugated drugs as the analyte. In the GMR assay, printed circuit boards (PCB) with eight fingers hold the GMR chips containing an array of 80 sensors per chip, which are spotted with the capture proteins. The capture reagent is spotted on the GMR chips at different concentrations to generate different loading densities. The sensor arrays are immersed in analyte solutions in a 96-deep-well microplate with vertical mixing (5 Hz) to minimize mass transport effects, as shown in **Figure 1A**. The PCB is moved from one row of reagent wells to the next to complete the association and dissociation phases of the assay. Unlike with the SPR platform, the analyte does not flow past the capture protein. **Figure 1B** compares the GMR and SPR measurements at the detection surface. In GMR, the MNP-analyte complex binding to the capture surface generates a magnetic field that causes a change in the electrical resistances of the underlying GMR sensors enabling the quantitative real-time monitoring of the binding kinetics. Only MNP-labeled molecules near the binding surface are detected. Direct binding from matrix components that are not labeled are not detected. In contrast, any molecule, analyte, or matrix component, that interacts with the SPR surface will generate a signal.

**Figure 1.**
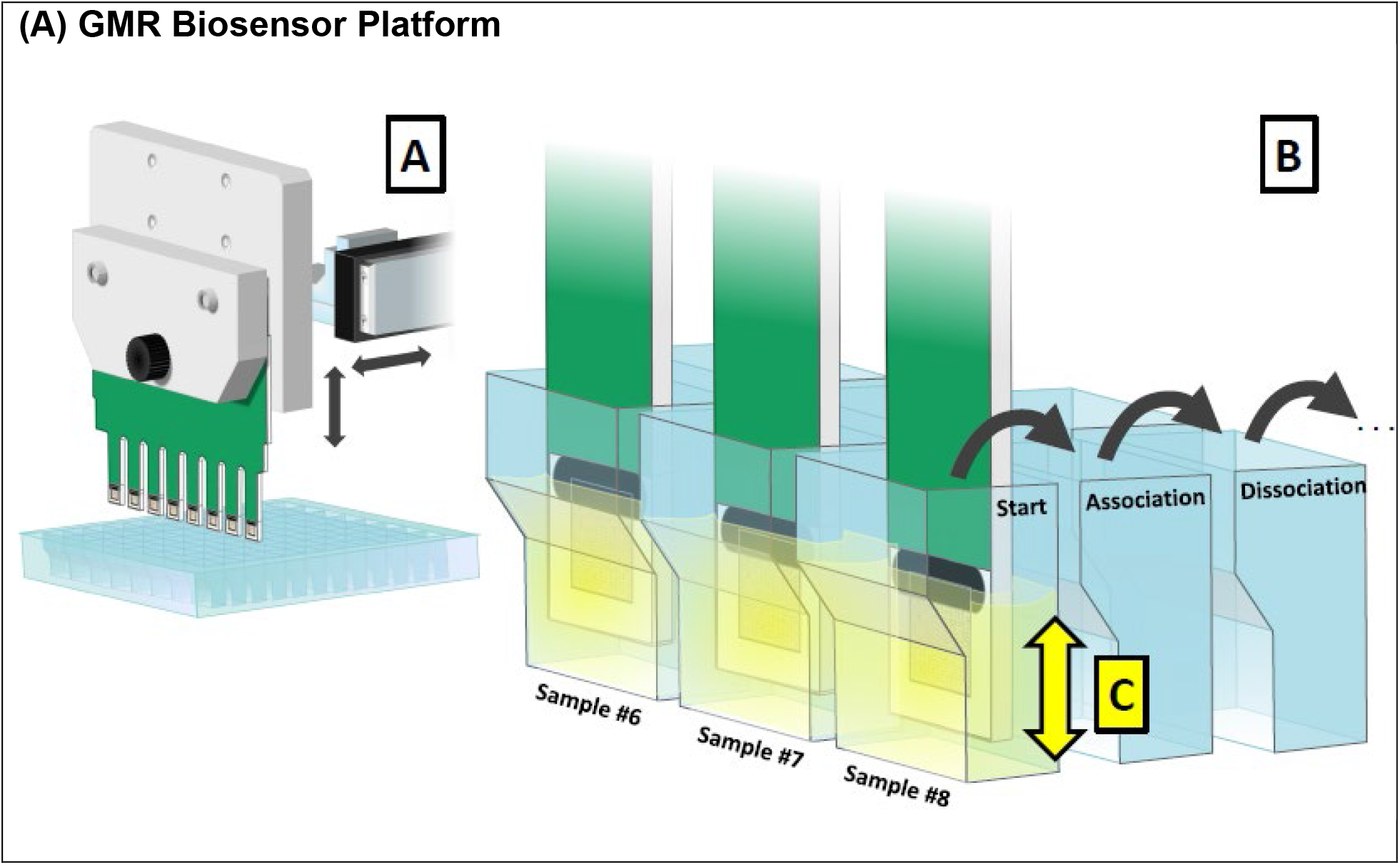
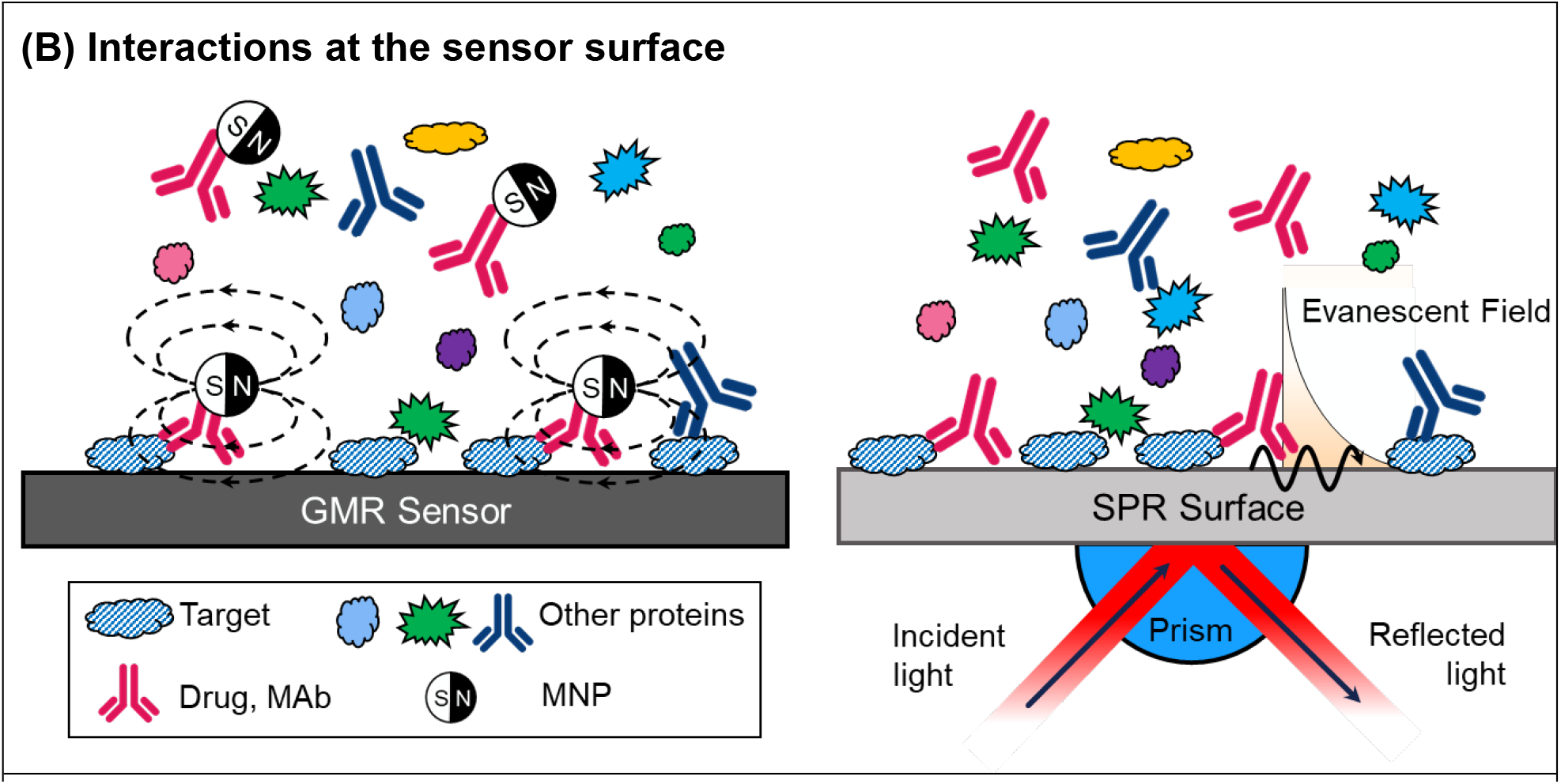
**(A) GMR biosensor platform. A**. A printed circuit board (PCB) with eight fingers extends into a custom 96 deep-well microtiter plate. Each PCB finger carries a chip with 80 GMR sensors that are individually printed with a drug, another ligand, or a reference protein and then immersed into the plate wells containing the MNP-drug conjugate. We typically spot 4-8 replicates of each capture probe together with multiple reference spots and average the results. Every sensor is continuously recorded at a signal integration time of 1 second. **B**. Each chip/well measures a separate ligand binding reaction, so that up to 8 ligand binding conditions can be evaluated in a single assay run. A robotic arm moves the PCB from one row of reagent wells to the next to complete the association and dissociation assay steps. **C**. The plate oscillates 2 mm vertically at 5 Hz to stir the reagents and minimize the formation of a local depletion zone around the sensor. **(B) Interactions at the sensor surface**. Comparison of the GMR and SPR measurements at the detection surface. In GMR, the MNP-analyte complex binding to the capture surface generates a magnetic field that causes a change in the electrical resistances of the underlying GMR sensor enabling the quantitative real-time monitoring of the binding kinetics. The capture protein on the chip and the assay conditions are established to favor one-to-one binding to avoid avidity complications. Only MNP-labeled molecules near the binding surface are detected. Labeled molecules that are away from the surface are not detected. Most importantly, direct binding of unlabeled matrix components is not detected. In contrast, any molecule, analyte, or matrix component, which interacts with the SPR surface, even weakly, will generate an SPR signal.

The binding curves for pembrolizumab with different PD-1 spotting concentrations are shown in **Figure 2A**. The MNP-drug conjugation ratio and concentration were optimized in separate experiments (data not shown). The sensorgrams are fitted with a mass transport model and the derived kinetic parameters are listed in **Table 1**. If a mass transport effect does not play a role in the pembrolizumab/PD-1 interaction, the k_on_ and k_off_ values will be independent of the surface density of PD-1. **Figure 2B** indeed shows there is no mass transport effect when the PD-1 spotting concentration is ≲14 mg/mL. This corresponds to a maximum DMR association signal of ∼920 PPM, indicating that the combination of lower spotting density, vertical mixing and an appropriate drug conjugate concentration, minimizes surface effects, notably, mass transport, rebinding and avidity, even for picomolar affinity interactions such as pembrolizumab/PD-1. One-to-one binding is being monitored under these conditions. Operationally, it is best to have a maximum ΔMR between ∼50 and ∼1000 PPM for kinetic assays. We have observed that this level minimizes surface effects and maintains a good signal-to-noise ratio.

**Figure 2.**
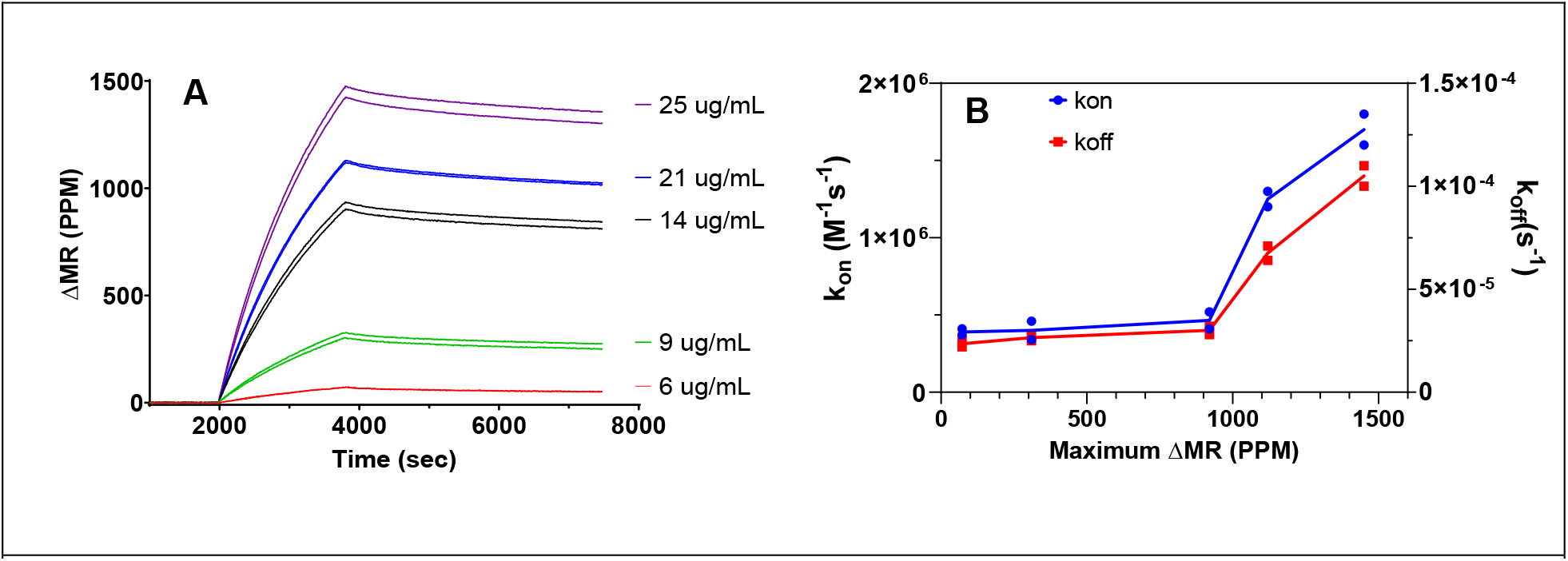
**(A) Pembrolizumab binding on a PD-1 spotted chip as a function of capture spotting concentration.** The sensorgrams are for the PD-1 capture reagent spotted on the chip at 6, 9, 14, 21 and 25 μg/mL. Binding curves are fit with a mass transport model and parameters are shown in **Table 1. (B) Association and dissociation rate constants for MNP-labeled pembrolizumab binding to PD-1 at different spotting concentrations**. Kinetic data from **Table 1** are plotted as a function of maximum ΔMR. The highest level of association (Max ΔMR) is the maximum value at each spotting concentration. The maximum ΔMR corresponds to spotting PD-1 at 6, 9, 14, 21 and 25 μg/mL, respectively. The binding constants begin to deviate at a ΔMR above 920 PPM, which corresponds to spotting at 14 μg/mL.

**Table 1.** GMR binding parameters for Pembrolizumab – PD-1 as a function of spotting concentration.

| Spotting ($\mu\text{g/mL}$) | Max $\Delta\text{MR}$ (PPM) | $k_{\text{on}}$ ($\times 10^5 \text{ M}^{-1} \text{ s}^{-1}$) | $k_{\text{off}}$ ($\times 10^{-5} \text{ s}^{-1}$) | $K_{\text{D}}$ (pM) |
| --- | --- | --- | --- | --- |
| 6 | 72 | $3.9 \pm 0.3$ | $2.4 \pm 0.2$ | $60 \pm 1.4$ |
| 9 | 310 | $4.0 \pm 0.8$ | $2.7 \pm 0.2$ | $68 \pm 9.2$ |
| 14 | 920 | $4.7 \pm 0.8$ | $3.0 \pm 0.3$ | $65 \pm 4.2$ |
| 21 | 1120 | $12.5 \pm 0.7$ | $6.8 \pm 0.5$ | $56 \pm 9.2$ |
| 25 | 1450 | $17.0 \pm 1.4$ | $10.5 \pm 0.7$ | $62 \pm 1.4$ |

### GMR, SPR and BLI biosensors give consistent results in buffer

Kinetic binding sensorgrams for the GMR, BLI and SPR platforms for three pairs of immune checkpoint inhibitors, atezolizumab/PD-L1, nivolumab/PD-1 and pembrolizumab/PD-1, are presented in **Figures 3A-C**, respectively. Results are shown for buffer with 0% serum and increasing concentrations of serum up to 80%. The kinetic binding parameters in buffer for the three platforms show modest agreement (**Table 2**). Using values from our measurements with the same proteins, the ratio of association rate constants of GMR to the other platforms is on average 2.2 and 1.6 for BLI and SPR, respectively. Dissociation rate constant ratios average 0.4 and 0.7 against the BLI and SPR platforms, respectively. K_D_ values calculated from the ratio of k_off_/k_on_ have an average ratio of 0.2 and 0.4, respectively. The Nivolumab /PD-1 pair tested in the reverse orientation, with nivolumab as the capture reagent, produced similar results with the PD-1 as the capture. It is widely recognized that it is difficult to harmonize kinetic measurements across platforms. Care is needed in making comparisons especially given the considerable discrepancies in the literature for the anti-PD-1 antibody binding constants, as indicated in **Table 2**.^15–17^ In general, it is best to compare drug-target interactions within the same platform using consistent experimental conditions.

**Figure 3.**
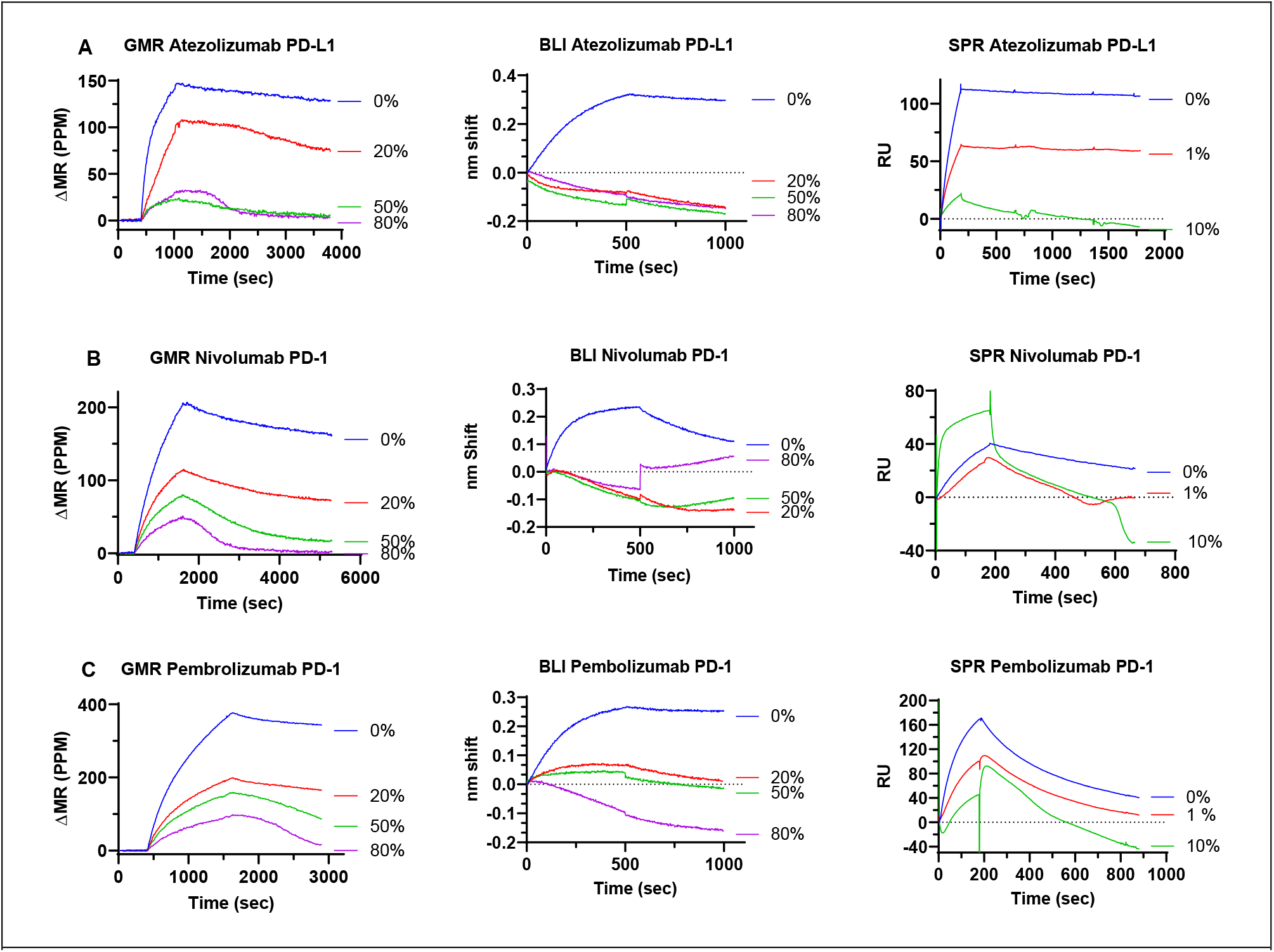
**(A**) Top row, binding of MNP labeled atezolizumab to PD-L1 with increasing concentrations of serum on the GMR, BLI and SPR platforms. GMR chip spotting concentration of PD-L1 is 8 μg/mL. Atezolizumab concentration are 4, 11 and 25 nM on the GMR, BLI and SPR platforms, respectively. **(B)** Middle row, binding of MNP labeled Nivolumab to PD-1 with increasing concentrations of serum on the GMR, BLI and SPR platforms. The GMR chip spotting concentration of PD-1 is 8 μg/mL. Nivolumab concentrations are 12, 33 and 25 nM on the GMR, BLI and SPR platforms, respectively. **(C)** Bottom row, binding of MNP labeled Pembrolizumab to PD-1 with increasing concentrations of serum on the GMR, BLI and SPR platforms. The GMR chip spotting concentration of PD-1 is 8 μg/mL. Pembrolizumab concentrations are 2, 11 and 25 nM on the GMR, BLI and SPR platforms, respectively.

**Table 2.** Binding constants in buffer.*

| Pair | Platform | $k_{on}$ ($\times 10^5$ M<sup>-1</sup>s<sup>-1</sup>) | $k_{off}$ ($\times 10^{-5}$ s<sup>-1</sup>) | $K_D$ (pM) |
| --- | --- | --- | --- | --- |
| Atezolizumab PD-L1 | GMR | 12 | 5.1 | 41 |
|  | GMR EQUIL | – | – | 43 |
|  | BLI | 4.0 | 12 | 277 |
|  | SPR | 5.7 | 3.1 | 54 |
| Nivolumab to PD-1 | GMR | 2.0 | 17 | 850 |
|  | GMR Rev** | 3.4 | 4.5 | 1400 |
|  | GMR EQUIL | – | – | 940 |
|  | BLI | 2.6 | 160 | 6200 |
|  | SPR | 2.3 | 44 | 1900 |
|  | SPRi-LSA<sup>15</sup> | 3.1 | 220 | 7200 |
|  | SPR TG<sup>18</sup> | 2.4 | 19 | 800 |
|  | SPR EMA<sup>16</sup> | 2.5 | 7.7 | 3100 |
| Pembrolizumab PD-1 | GMR | 11 | 6.1 | 56 |
|  | GMR EQUIL | – | – | 101 |
|  | BLI | 4.0 | 12 | 280 |
|  | SPR | 5.7 | 220 | 3900 |
|  | SPRi-LSA<sup>15</sup> | 6.7 | 270 | 3900 |
|  | SPR TG<sup>18</sup> | 4.2 | 140 | 3400 |
|  | SPR EMA<sup>17</sup> | 10.4 | 3.1 | 29 |

\*Results are based on the curves for buffer (0% serum) in **Figure 3 and cited references**.<sup>15-18</sup>
\*\*GMR kinetic assay with nivolumab as the capture and MNP-PD-1 as the analyte.

We also evaluated Nivolumab and Pembrolizumab binding to PD-1 with an indirect labeling approach where the drugs were first biotinylated and then mixed with 20-nm streptavidin (SA) coated MNP (see **Supplemental Method)**. The binding measurements were then carried out as with the direct MNP label approach. The k_on_, k_off_ and K_D_ values for both Nivolumab/PD-1 and Pembrolizumab/PD-1 are in good agreement between the two methods (**Supplemental Table 2**). We also successfully applied this indirect labeling method to five additional protein pairs: PD-1/PD-L1, Bevacizumab/VEGF, Adalimumab/TNFα, Infliximab/TNFα, and Axl/Gas6. The results demonstrate that the GMR kinetic platform can be applied to determine a broad range of binding constants. Association rate constants ranged 170-fold, from 0.3 to 50 ×10^5^ M^-1^s^-1^. Dissociation rate constants ranged 150-fold, from 0.15 to 230 ×10^−4^ s^-1^ and calculated K_D_ values ranged more than 40,000-fold, from 18 to 800,000 pM. In agreement with the SPR literature^19^, the fasted off-rate and lowest affinity was for the PD-1/PD-L1 pair(**Supplemental Table 2**).

We further evaluated the binding affinities for the Nivolumab/PD-1, Pembrolizumab/PD-1 and Atezolizumab/PD-L1 with an equilibrium binding assay. Titration curves for binding on the GMR platform configured as a sandwich immunoassay^8^ are shown in **Figure 4**. K_D_ values, calculated by five-parameter logistic regression (5PL) are included in **Table 3** for the 0% serum matrix. The GMR equilibrium results are in good agreement with the GMR kinetic results, with differences of 5, 11 and 80% for the 3 pairs. The cross-platform kinetic comparisons and the within-platform kinetic and equilibrium comparisons strongly validate the GMR kinetic platform. Kinetic measurements can be successfully made on the GMR platform with magnetic bead conjugated analytes and optimized assay conditions.

**Figure 4.**
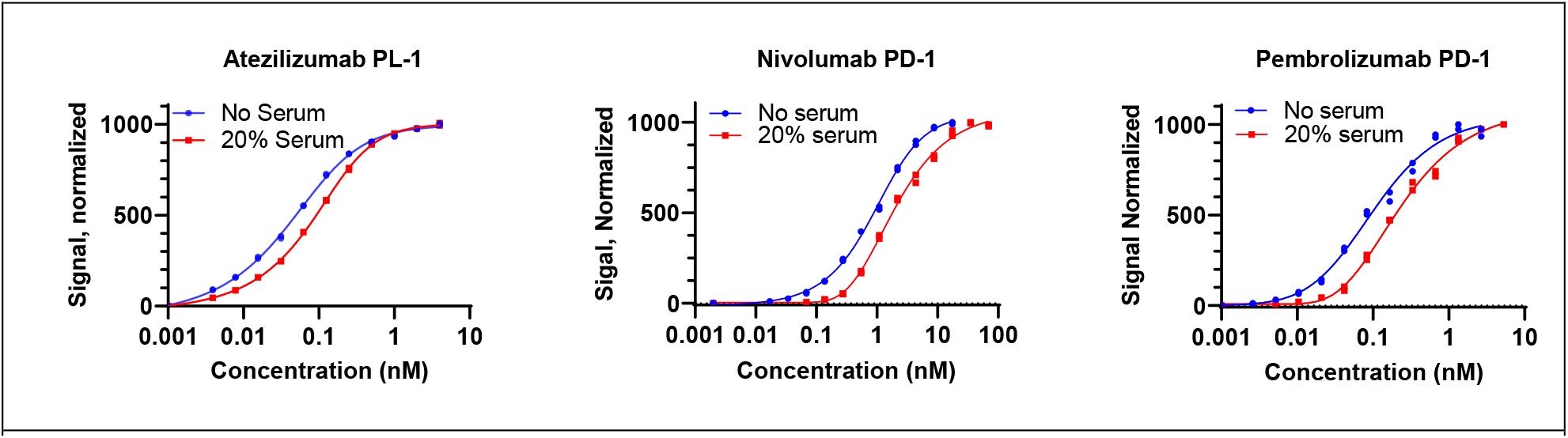
K_D_ evaluation with an equilibrium GMR assay. Full titration curves for Atezolizumab/PD-L1, Nivolumab/PD-1 and Pembrolizumab/PD-1 at 0 and 20% serum. Data are normalized for easier visualization. Solid lines are the 5PL curve fit used to calculate K_D_. All 6 curves had a goodness of fit R^2^ > 0.99.

**Table 3.**
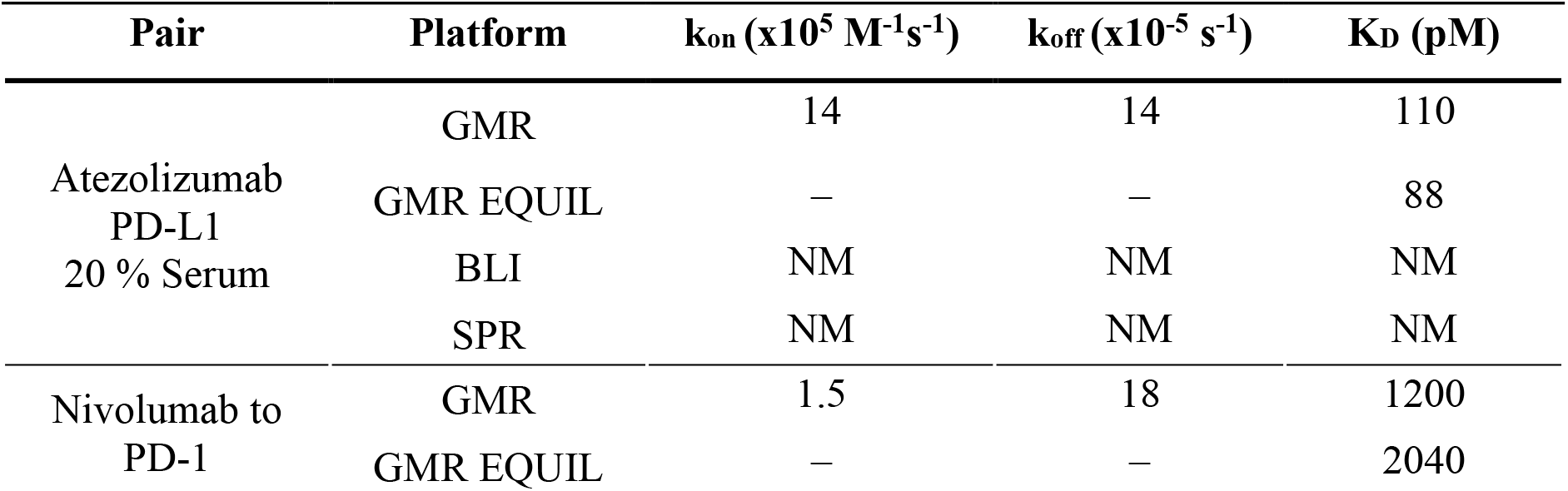
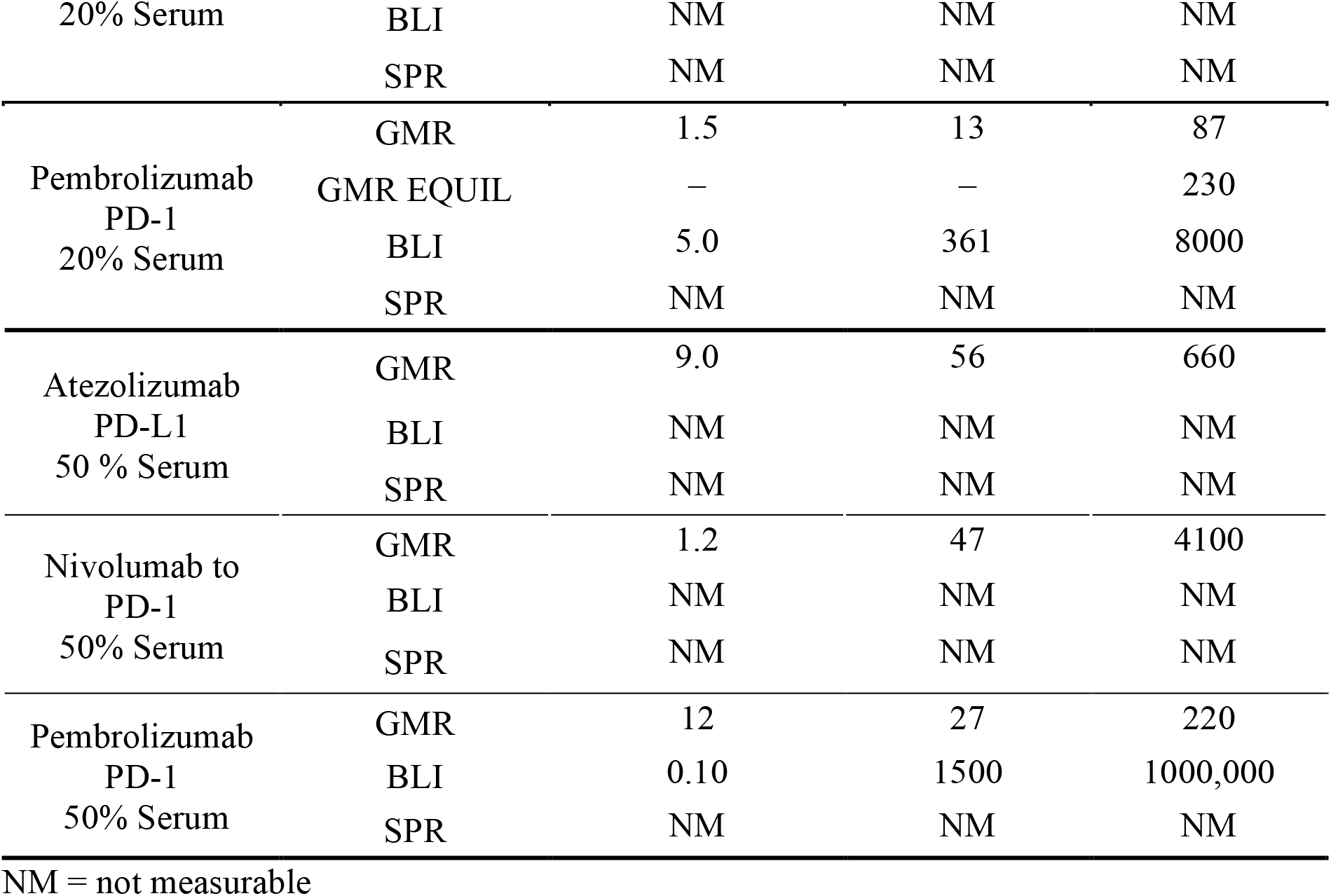
Binding constants in serum.

### The GMR biosensor platform is best suited for determining binding kinetic parameters with increasing amounts of serum

Kinetic binding sensorgrams with varying levels of serum for the three pairs of immune checkpoint inhibitors were shown earlier in **Figure 3** for all three platforms. Visual inspection of the curves readily shows the platform differences. The SPR and BLI platforms have severe limitations due to the serum matrix, likely an interference with the optical signals and cannot distinguish between specific binding and non-specific binding of serum proteins.^10,20,21^ As a result, negative signals are sometimes observed after reference subtraction, or the derived kinetics parameters are not reliable, even if the binding traces can be fit. The SPR platform is not able to generate suitable curves in as little as 10% serum. The BLI platform also has great difficulty generating curves suitable for binding constant evaluation in 20 and 50% serum. In striking contrast, the GMR platform can generate quality sensorgrams in up to 80% serum for all three drug-target pairs. Kinetic parameters could be derived with the GMR platform with 0, 20 and 50% serum for all three pairs, using nonlinear regression analysis of the primary sensorgram data with simple 1-to-1 or mass transport binding models (**Figure 5** and **Table 3**).

**Figure 5.**
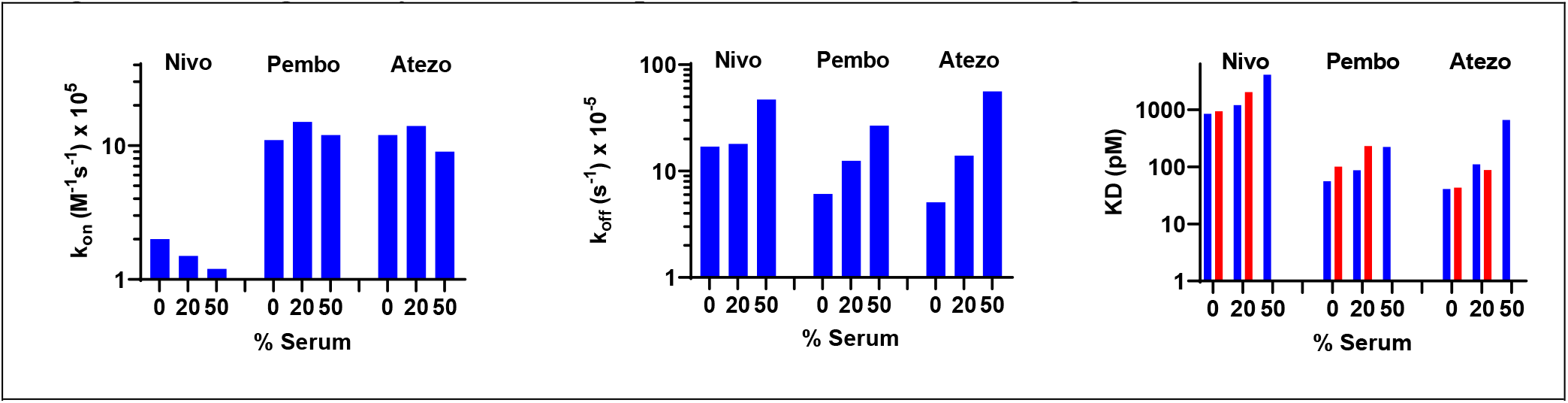
GMR-derived binding constants for immune checkpoint inhibitor pairs with increasing serum concentrations. Values for k_on_, k_off_ and K_D_ are graphed for nivolumab/PD-1, pembrolizumab/PD-1 and atezolizumab/PD-L1 on a log scale. 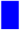 Kinetic assay, 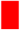 Equilibrium assay.

The GMR derived k_on_ values are relatively unaffected by increasing concentrations of serum. The values are within 10 to 40% in the presence of 20 or 50% serum for the three drug-target pairs. Only MNP-labeled molecules are detected in the GMR kinetic assay where binding from un-labeled matrix components does not interfere. These results indicate there is little or no effect of albumin or other serum components on the rates of association for these three drug-target pairs.

In contrast, k_off_ values are generally higher with increasing concentrations of serum for these three drug-target pairs. At 50% serum, the k_off_ values are 2.8, 4.4 and 11-fold higher for nivolumab, pembrolizumab and atezolizumab, respectively. This suggests specific or nonspecific interactions with serum components affects the k_off_ rates more than the k_on_ rates for these the drug-target pairs. This may be different for other protein-protein pairs. Consistent with the higher k_off_ rates and similar k_on_ rates, the K_D_ values (as k_off_/ k_on_) are higher with increasing amounts of serum. At 50% serum, the K_D_ values are 4.8, 4.0 and 16-fold higher for nivolumab, pembrolizumab and atezolizumab, respectively. For both k_off_ and K_D_, the serum effect was greater for the PDL-1 drug, atezolizumab, than the two PD-1 drugs, nivolumab and pembrolizumab. The kinetics of bevacizumab binding to VEGF are also strongly affected by serum. At 50% serum, both k_off_ and K_D_ are 20-fold higher **(Supplemental Figure 2 and Supplemental Table 2**). We validated the accuracy of the GMR kinetic K_D_ results with the GMR equilibrium assay results in both 0% and 20% serum (**Figure 5, Table 4**). The two methods are in good agreement for all three checkpoint inhibitor antibody pairs.

The higher k_off_ values and hence shorter drug-target residence times, suggest specific interactions between serum components and drug-target pairs that weaken the binding. This faster off rate could be due to several factors such as protein conformational changes for the receptor ligand pair, or a greater binding affinity for the serum proteins rather than for the ligand.

## Discussion

It is increasingly appreciated that the rates at which molecules associate with and dissociate from receptors — the binding kinetics — directly impact the therapeutic index of an effector molecule.^22^ Since being introduced in 2006, drug-target residence time models have been widely accepted.^23–26^ However, binding affinity, K_D_, alone is insufficient in determining pharmacological activity. Rather, it is the residence time, τ = 1/k_off_, of the drug-target complex that determines much of the *in vivo* pharmacological activity. Most early studies of drug activity were carried out in closed *in vitro* systems where the drug molecule was present at an unchanging concentration. In contrast, open *in vivo* systems experience constantly changing local drug concentrations because of multiple physiological processes such as absorption, tissue distribution, metabolism and excretion (ADME). Binding kinetics are therefore influenced by factors other than just the molecular determinants of receptor-ligand interactions. Specific and non-specific binding by serum proteins can influence the binding behavior of receptors and ligands. The introduction of the SPR technology led to the ability to systematically consider association and dissociation kinetics. However, that platform is limited in its ability to make measurements in physiologically relevant matrices.

GMR biosensors are proximity-based detectors of dipole fields from the magnetic tag, so only MNP-labeled molecules within ∼150 nm of the surface are detected.^8,9^ Unbound MNP-molecules and unlabeled molecules are not detected, making the platform ideal for real-time kinetic analysis. This is an important advantage compared to SPR and BLI techniques where assignment of interacting molecules is very difficult because analyte and non-analyte molecules cannot be distinguished. Kinetic information that includes the influence of the biological matrix may allow more reliable conclusions to be made on protein functionality and facilitate more efficient drug development.

The comparisons of the three immune checkpoint inhibitors in the presence and absence of serum are striking. While the k_on_ values remained relatively constant, the k_off_ values are higher with increasing concentrations of serum for these three drug-target pairs. At 50% serum, the k_off_ values are 2.8, 4.4 and 11-fold higher for nivolumab, pembrolizumab and atezolizumab, respectively. Previously, our colleagues used the GMR platform to demonstrate that even small amounts of a serum-based product had a profound effect on the binding of protein kinase A with its small molecule target quercetin.^10^ In those experiments, the serum-based product^27^, had little effect on two antibody drug-target pairs, infliximab/TNFα and bevacizumab/VEGF. Thus, the processed serum-based product may not be an accurate substitute for subject serum. In a limited set of studies, we see similar kinetic results with serum samples from different single healthy donors (data not shown). It would be worthwhile to evaluate the effects of serum samples from different induvial donors with different clinical indications and with different drug treatments in a follow-up study.

The kinetic binding properties of immune checkpoint inhibitors should be considered as part of the clinical application of the drugs. Our results show the biggest serum effect in both k_off_ and K_D_ for the PDL-1 drug, atezolizumab, compared with the two PD-1 drugs, nivolumab and pembrolizumab. The effect on the PD-1 drugs is more subtle. Much of the clinical data suggest that the two anti-PD-1 inhibitors, nivolumab and pembrolizumab are interchangeable. They are approved for a partially overlapping but distinct set of indications. Such differences may be due to either drug-dependent or drug-independent reasons.^28^ Pembrolizumab and nivolumab have significant molecular similarities.^29,30^ Both drugs target epitopes on the PD-1 molecule with high affinity and specificity. Both are IgG4 and hence have similar effector functions which are minimal for this subclass. They have essentially identical amino acid sequences except for the variable regions that bind different epitopes of the antigen.^28^ Their receptor occupancy saturation is the same and their pharmacokinetic properties, such as clearance, terminal half-life, steady state concentrations and recommended dosages, are indistinguishable.^31^ Yet, molecular studies show differences between nivolumab and pembrolizumab. Of particular interest is the elucidation of the PD-1/pembrolizumab-FAb complex structure comparing the binding modes and blockade mechanisms of nivolumab and pembrolizumab.^32^ These two antibodies showed a similar binding mode to PD-1, in which the dominant interaction was located on the loops of PD-1 and the competitive binding with PD-L1 involved a steric clash that prevented the binding of PD-1 to PD-L1. However, the targeted loops are completely different: pembrolizumab mainly binds to the CD loop, whereas nivolumab mainly binds to the N-loop, with no overlapping binding areas on PD-1, or with each other.^31^ These differences may be important in considering the effects of serum on the dissociation rates. Such serum effects have been noted before, for example, in the binding differences of the anti-cocaine antibody binding to cocaine and its metabolites in buffer compared to serum.^33^ Since different binding kinetics have been known to cause different physiological responses in different therapeutic indications for two drugs binding to the same target, we suggest that a similar mechanism might account for the differential clinical outcomes of nivolumab and pembrolizumab. Furthermore, it may be useful to determine the impact of serum samples from individuals with different clinical indications and different therapeutic treatments. Such measurements may provide insight into primary and acquired resistance to immune checkpoint blockade therapies.^34^ The high sensitivity and matrix-insensitivity of the GMR kinetic assay platform will be helpful in elucidating new drug targets for immunotherapy.^33-34^

Given the profound impact of biologics in cancer immunotherapy and the growing number of immune checkpoint inhibitors^11–13^, it is of increasing importance to understand how interchangeable monoclonal antibody inhibitors are likely to be when they share a common therapeutic target. Our results suggest the ability to measure binding kinetics in a serum matrix from different subjects is important to that understanding.

In summary, we have established GMR biosensors as a capable platform technology for the real-time monitoring of complicated molecular interactions in biologically relevant matrices. Its adoption for evaluating the differential binding kinetics of drug-target pairs in serum will be an important paradigm shift for the field. Understanding the molecular interactions that occur in a biological matrix is indispensable for determining the mechanism of action of drugs and the pharmacokinetics/ pharmacodynamics inside the body.

## Methods

### MNP-protein conjugation

To generate MNP complexes for kinetic assays, nivolumab (BioVision, #A1307), pembrolizumab (BioVision, #A1306 and atezolizumab (BioVision, #A1305) were individually mixed with NHS-activated MNPs (Miltenyi Biotec, #130-105-805) at a molar ratio of 10:1 and incubated at 4°C overnight with gentle mixing. After the incubation, the conjugated MNPs were purified with a magnetic micro-column (Miltenyi Biotec, #130–042-701). Briefly, the mixture was added to a column placed under a strong magnetic field and the MNPs were trapped on the column. After washing away unbound proteins, the external magnetic field was removed and the MNPs were eluted with PBS containing 1% BSA and 0.05% Tween-20.

### GMR platform

The MagArray MR-813 12R platform (Milpitas, CA) was used for GMR measurements. The sensing board consisted of 8 chips per board and 80 sensors per chip (**Figure 1A**). To create multiple loading densities a series of concentrations of PD-1 (R&D systems, #8986-PD) or PD-L1 (R&D systems, #156-B7) were applied to the sensors using a spotter (Scienion, S5). For the reference sensors, 0.1% bovine serum albumin (BSA) solution was spotted. After spotting, the chips were blocked with a protein blocking solution. The magneto-resistance measurements are carried out with the MagArray MR-813 12R reader system. The reagents were placed in a 96-deep well plate (MagArray) contained within the instrument and the position of the GMR sensor was automatically controlled inside and outside of each deep well where sample or kinetic buffer were placed. The ratio of the resistance of the GMR sensor before and after dipping the GMR sensor into the MNP dispersed solution was calculated as ΔMR. To determine the baseline signal (for initial sensor offset zeroing), the target protein spotted GMR sensor chips were dipped into kinetic buffer (PBS with 1% BSA and 0.05% Tween-20), or into kinetic buffer containing 20, 50 or 80% single donor human serum (Golden West BioSolutions, Temecula, CA) for 1 min. For the association reaction process, the GMR sensor chips were dipped into wells containing the MNP-drug conjugate solution for 30 min and the signal from the GMR sensor was monitored in real-time with one data point acquisition every 5s. The GMR sensor chips were then dipped into wells containing only kinetic buffer and the dissociation signal was monitored. The kinetic buffer used in the baseline, association and dissociation steps were matched. Vertical mixing (2 mm, 5 Hz) was used to minimize mass transport effects.

The Scrubber software package (Biologic Software, Campbell Australia) was used for the GMR curve fitting. For nivolumab, the dissociation (k_off_) and association (k_on_) rate constants were obtained by nonlinear regression analysis of the primary sensorgram data according to a 1:1 binding model. For pembrolizumab and atezolizumab, the mass transport fitting model was used. The dissociation constant K_D_ was calculated using the formula K_D_ = k_off_ /k_on_.^35^

### SPR platform

The SPR kinetic interactions of the three proteins were measured by Yurogen Biosystems LLC (Worcester, MA) at 25°C using a Biacore 3000 biosensor (Biacore AB, Uppsala, Sweden) as described previously^14^. Drug and target materials supplied to the vendor were the same as for the GMR measurements. After immobilization of the antibodies onto the carboxymethylated dextran surface of a CM5 sensor chip at a level of 1000-2000 response units, 60 µL of kinetic buffer containing 0, 1 or 10 % serum was injected into the flow cell at a rate of 30 µL/min. As a control, BSA (0.04 mg/mL) was simultaneously immobilized onto the reference surface under the same conditions to correct for instrument and buffer artifacts.

### BLI platform

The BLI kinetic assays were performed by Antibody Solutions (Santa Clara, CA) using an Octet RED biosensor (ForteBio, Fremont, CA) by first capturing the monoclonal antibody (MAb) using anti-human Fc (AHC) biosensors, followed by a baseline step of 300 s in kinetic buffer or kinetic buffer containing 20, 50, or 80% human serum. Drug and target materials supplied to Antibody Solutions were the same as for the GMR measurements. The MAb-captured biosensors were then submerged into wells containing different concentrations of antigen for 500 s followed by 500 s of dissociation in the kinetic buffer. The kinetic buffer used in the baseline, association and dissociation process matched with one another. The MAb-captured sensors were also dipped in kinetic buffer or kinetic buffer containing 20, 50, or 80% serum to allow single reference subtraction to compensate for the natural dissociation of captured MAb. The binding sensorgrams were collected using the 8-channel detection mode on the Octet RED96 unit. Fresh AHC biosensors were used without any regeneration step.

### Equilibrium Assays

Equilibrium assays were performed with the MagArray MR-813 reader system. The capture antibody pembrolizumab was spotted on the GMR sensor surface at the concentration of 100 µg/mL. Next, the GMR sensors were allowed to incubate with serial dilutions of biotinylated PD-1 prepared in buffer or buffer plus 20% human serum for 3 hours. After a washing step, the sensors were then dipped into a solution containing anti-biotin antibody coated magnetic beads (Miltenyi) with constant mixing to form an immunoassay complex. Data were recorded at the signal maximum after 20 minutes and fit with five-parameter logistic regression (5PL) using GraphPad Prism version 8.0.0 for Windows (GraphPad Software, San Diego, CA). K_D_ values were calculated as the C_50_ concentrations of the 5PL curve.

## Acknowledgements

We thank Mike Beggs for thoughtful review of the manuscript. We thank Adam Seger for contributions to **Figure 1B**. All data needed to evaluate the conclusions in the paper are present in the paper and/or the Supplementary Materials. The authors acknowledge that they received no funding in support for this research

## Author Contributions

DY contributed to the experimental design, conducted the experiments, analyzed data and contributed to the writing of the manuscript. HY and TS provided technical advice on the GMR platform. HY and SJO engineered the GMR biosensor platform for the kinetic assays. ABK contributed to the experimental design, data analysis, data interpretation and wrote the main manuscript with input from other authors. SXW contributed to the design and scientific content of the project. KC initiated the project and contributed to the experimental design, data analysis, data interpretation and writing of the manuscript. KC, MLC and SXW supervised the project.

## Corresponding Authors

Please direct all correspondence to Prof. Shan Wang or Dr. Kalidip Choudhury.

## Supplementary Materials

Binding kinetics can help differentiate therapeutic properties for drugs that bind the same target.^1^ For example, the calcium channel blocker verapamil, which has a slower rate of dissociation, is used to treat supraventricular tachyarrhythmias, while nifedipine with a faster dissociation rate is used to primarily treat hypertension.^2^ Similarly, occupancy profiles for the angiotensin II receptor inhibitors losartan and candesartan could not explain the increase in efficacy observed with candesartan which displays slow dissociation kinetics.^3^ In both these examples the differential binding kinetics correlate with the differential clinical response.

**Supplementary Table 1.** Mechanistic differentiation of medicines that bind to the same target protein.

| Medicines | Target | Differential Clinical Outcome | Biochemical differentiation |
| --- | --- | --- | --- |
| **Candesartan**<br>**Losartan** | Angiotensin II receptor (GPCR) | Maximum efficacy | Slow dissociation kinetics |
| **Clozapine**<br>**Haloperidol** | Dopamine receptor (GPCR) | Improved safety | Fast kinetics |
| **Verapamil**<br>**Dihydropyridine** | Ca<sup>+</sup> ion channel | Therapeutic indication | kinetics |
| **Lidocaine**<br>**Lecainide** | Na<sup>+</sup> ion channel | Therapeutic indication | kinetics |
| **Estrogen**<br>**aloxifene** | Estrogen receptor | Safety | Unique conformational state |

After Swinney<sup>1</sup>

**Supplementary Table 2.**
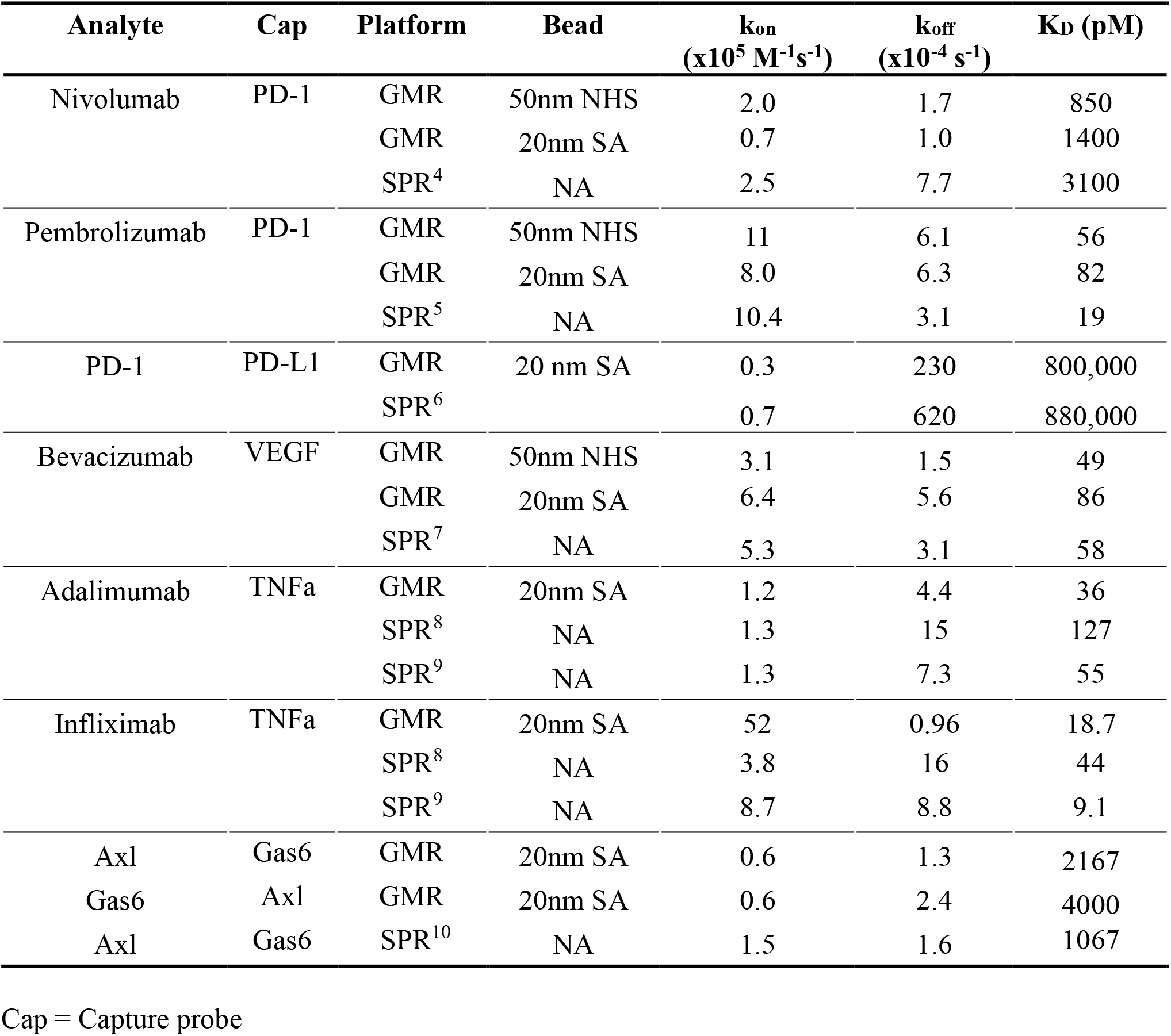
Binding constants measured by GMR biosensor and SPR platforms.^4-10^

| Analyte | Cap | Platform | Bead | $k_{on}$<br>($\times 10^5 \text{ M}^{-1} \text{ s}^{-1}$) | $k_{off}$<br>($\times 10^{-4} \text{ s}^{-1}$) | $K_D$ (pM) |
| --- | --- | --- | --- | --- | --- | --- |
| Nivolumab | PD-1 | GMR | 50nm NHS | 2.0 | 1.7 | 850 |
|  |  | GMR | 20nm SA | 0.7 | 1.0 | 1400 |
|  |  | SPR<sup>4</sup> | NA | 2.5 | 7.7 | 3100 |
| Pembrolizumab | PD-1 | GMR | 50nm NHS | 11 | 6.1 | 56 |
|  |  | GMR | 20nm SA | 8.0 | 6.3 | 82 |
|  |  | SPR<sup>5</sup> | NA | 10.4 | 3.1 | 19 |
| PD-1 | PD-L1 | GMR | 20 nm SA | 0.3 | 230 | 800,000 |
|  |  | SPR<sup>6</sup> |  | 0.7 | 620 | 880,000 |
| Bevacizumab | VEGF | GMR | 50nm NHS | 3.1 | 1.5 | 49 |
|  |  | GMR | 20nm SA | 6.4 | 5.6 | 86 |
|  |  | SPR<sup>7</sup> | NA | 5.3 | 3.1 | 58 |
| Adalimumab | TNFa | GMR | 20nm SA | 1.2 | 4.4 | 36 |
|  |  | SPR<sup>8</sup> | NA | 1.3 | 15 | 127 |
|  |  | SPR<sup>9</sup> | NA | 1.3 | 7.3 | 55 |
| Infliximab | TNFa | GMR | 20nm SA | 52 | 0.96 | 18.7 |
|  |  | SPR<sup>8</sup> | NA | 3.8 | 16 | 44 |
|  |  | SPR<sup>9</sup> | NA | 8.7 | 8.8 | 9.1 |
| Axl | Gas6 | GMR | 20nm SA | 0.6 | 1.3 | 2167 |
| Gas6 | Axl | GMR | 20nm SA | 0.6 | 2.4 | 4000 |
| Axl | Gas6 | SPR<sup>10</sup> | NA | 1.5 | 1.6 | 1067 |

Cap = Capture probe

**Supplementary Figure 1.**
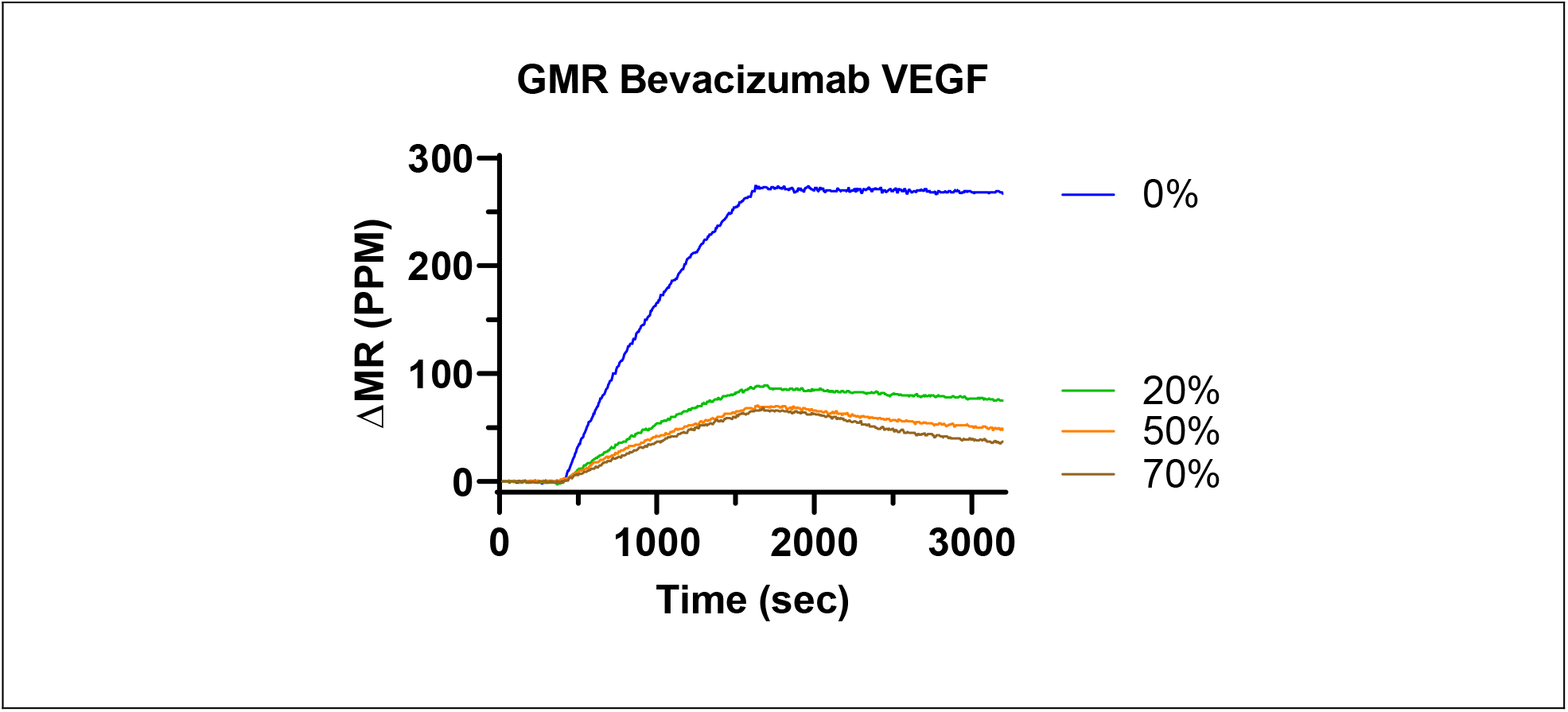
Binding of bevacizumab to VEFG with increasing concentrations of serum, measured with the GMR biosensor platform.

## Supplemental Method

**Supplemental Table 2** contains GMR biosensor results with two different assay methods. The first approach is the same as in the main body of the paper, where the analyte proteins are individually conjugated to 50-nm NHS-activated magnetic beads from Miltenyi Biotec. In the second approach, the proteins are first individually biotinylated with NHS-PEG_4_-Biotin (ThermoFisher Scientific, #A39259) and purified using desalting columns (ThermoFisher Scientific, #89890). The biotinylated proteins are then each mixed with 20-nm streptavidin (SA) coated MNP (Ocean Nanotech, San Diego, CA, #SHS-20-5) at a 3:1 molar ratio with gentle mixing for 15 min. The conjugated MNPs are then purified with a magnetic micro-column (Miltenyi Biotec, #130–042-701). Proteins from BioVision include nivolumab (#A1307) pembrolizumab (#A1306, Atezolizumab (#A1305), Adalimumab (# A1048), Infliximab (#A1097), PD-L1 (#8986-PD), PD-1 (#156-B7), TNFa (# 10291-TA-050) Gas6 (#885-GSB-050) and Axl (#10448-AL-100). Bevacizumab (#HY-P9906) is from MedChemExpress and VEGF is from Peprotech (#100-20).

**Supplementary Table 2.** Additional binding results in serum and buffer and the ratios of kinetic parameters in serum (20% and 50%) over those in buffer (0% serum).

| Analyte | Cap | Platform | % Serum | $k_{on}$<br>($\times 10^5 \text{ M}^{-1}\text{s}^{-1}$) | $k_{off}$<br>($\times 10^{-5} \text{ s}^{-1}$) | $K_D$<br>(pM) |
| --- | --- | --- | --- | --- | --- | --- |
| Bevacizumab | VEGF | GMR | 0 | 4.2 | 1.2 | 29 |
|  |  | SPR<sup>7</sup> | 0 | 5.3 | 3.1 | 58 |
|  |  | GMR | 20 | 3.7 | 9.5 | 256 |
|  |  | GMR | 50 | 3.7 | 23 | 630 |
| Ratio |  | GMR | 20%/ 0% | 0.88 | 7.9 | 9.0 |
| Ratio |  | GMR | 50%/ 0% | 0.88 | 20 | 22 |

